# Mutational processes of distinct POLE exonuclease domain mutants drive an enrichment of a specific TP53 mutation in colorectal cancer

**DOI:** 10.1101/681767

**Authors:** Hu Fang, Jayne A. Barbour, Rebecca C. Poulos, Riku Katainen, Lauri A. Aaltonen, Jason W. H. Wong

## Abstract

Cancer genomes with mutations in the exonuclease domain of Polymerase Epsilon (POLE) present with an extraordinarily high somatic mutation burden. *In vitro* studies have shown that distinct POLE mutants exhibit different polymerase activity and yet, how these POLE mutants generate mutations across cancer genomes and influence driver events remains poorly understood. Here we analyzed 7,345 colorectal cancer samples, including nine whole genome sequenced samples harboring *POLE* mutations. Our analysis identified differential mutation spectra across the mutants including methylation-independent enrichment of C>T mutations in POLE V411L. In contrast, analysis of other genomic regions showed similar mutation profiles across the different POLE mutants. Notably, we found that POLE mutants with the TP53 R213* mutation, caused by a TT[C>T]GA substitution, have significantly higher relative frequency of this mutational context compared with samples without this mutation. This finding demonstrates that variations in underlying mutation spectra can increase the likelihood of specific driver mutation formation.

## Introduction

*POLE* encodes the catalytic subunit of DNA Polymerase Epsilon, which is responsible for DNA fidelity during the process of eukaryotic nuclear genome replication (Jansen et al., 2016). Functional POLE mutations have been identified in less than 1% of all cancer genomes but these genomes are characterized by exceptionally high tumor mutation burden (Campbell et al., 2017). Somatic mutations of POLE exonuclease domain are frequently enriched in brain, uterine and colorectal cancer, and patients with POLE dysfunction usually have significantly better prognosis and require less intensive treatment (Cancer Genome Atlas Research et al., 2013).

The POLE mutational process shapes the cancer genome into a unique mutational signature with high proportions of C>A mutations at TCT contexts, C>T mutations at TCG contexts and T>G mutations at TTT contexts, which is known as COSMIC signature 10 (Alexandrov et al., 2013a). Several driver mutations have been identified in the *POLE* exonuclease domain (codons 268–471) (Church et al., 2013), the most frequent being P286R and V411L (Campbell et al., 2017). Residue P286 is located in the DNA binding pocket which is adjacent to the exonuclease active site. A change of this amino acid has been predicted to affect the structure of the DNA binding pocket and cause polymerase hyperactivity (Xing et al., 2019). By contrast, residue V411 lies a distance away from the binding site and does not interact with the DNA sequence directly (Briggs and Tomlinson, 2013). Data showing the mutation spectrum of individual POLE mutants supports differences in the way mutants generate somatic mutations (Shinbrot et al., 2014) but these differences have not yet been quantified.

Mutations are distributed unevenly across the cancer genome and mutation rates across genomic regions are highly heterogeneous (Gonzalez-Perez et al., 2019) due to due to genomic and epigenetic features including cytosine methylation (Walsh and Xu, 2006), replication timing (Koren et al., 2012), tri-nucleotide/penta-nucleotide context composition (Alexandrov et al., 2013a), transcription factor binding, chromatin organization (Schuster-Böckler and Lehner, 2012), gene expression levels (Hodgkinson and Eyre-Walker, 2011), orientation of the DNA minor groove around nucleosomes (Pich et al., 2018), CTCF binding (Poulos et al., 2016) and gene body features such as introns and exons (Frigola et al., 2017). As POLE mutant cancers are usually hypermutated and individual mutants might lead to distinct mutator phenotypes, the precise mechanisms of mutagenesis may be revealed by investigating whether they show disparity in mutational spectrum and distribution across genomic regions.

In this study, we first characterized 53 whole genomes of colorectal cancer, which harbor different *POLE* exonuclease domain somatic mutations (n=9) or are POLE wild-type (n=44). The mutational spectrum of the different POLE mutants was compared and validated in additional whole exome/target capture sequencing colorectal cancer samples. We also studied the association between cytosine methylation and mutation burden, and examined genome-wide mutation profiles across a range of genomics features. Finally, combining these datasets, we sought to identify associations between specific POLE mutants and the formation of driver mutation hotspots in colorectal cancer.

## Results

### Profile of mutation signatures in different POLE-mutant colorectal cancers

A total of 53 colorectal cancer whole genomes were analyzed, in which 44 are POLE wild-type and microsatellite stable, and the remaining nine carried non-synonymous somatic mutations in the POLE exonuclease domain (**Supplementary table 1**). We clustered these samples based on the proportion of 96 tri-nucleotide contexts and obtained four distinct groups (**Figure 1A**). The nine POLE mutants were clustered into three subgroups, which are represented as P286R (n=3), V411L (n=3) and Other-Exo (n=3) (sample with other mutations in the POLE exonuclease domain). In line with previous reports (Shinbrot et al., 2014), all of the POLE mutants showed a high proportion of C>A and T>G mutation in TCT and TTT tri-nucleotide contexts, which resembles COSMIC signature 10 (**Figure 1B**). When examining genome-wide C>T mutations, we observed a higher proportion of C to T mutations in POLE V411L mutants accounting for 33.7% of all substitutions, while there were 16.0% and 23.3% in P286R and Other-Exo mutants respectively (P286R vs V411L, P<0.001, Chi-squared test, **Figure 1C**). The differential mutation spectrum clustering of P286R from V411L mutants was also evident in 32 POLE mutant colorectal cancer samples with WGS, WXS and target capture sequencing data (**Figure 1-figure supplement 1**), confirming the differences observed in the WGS samples.

**Figure 1.**
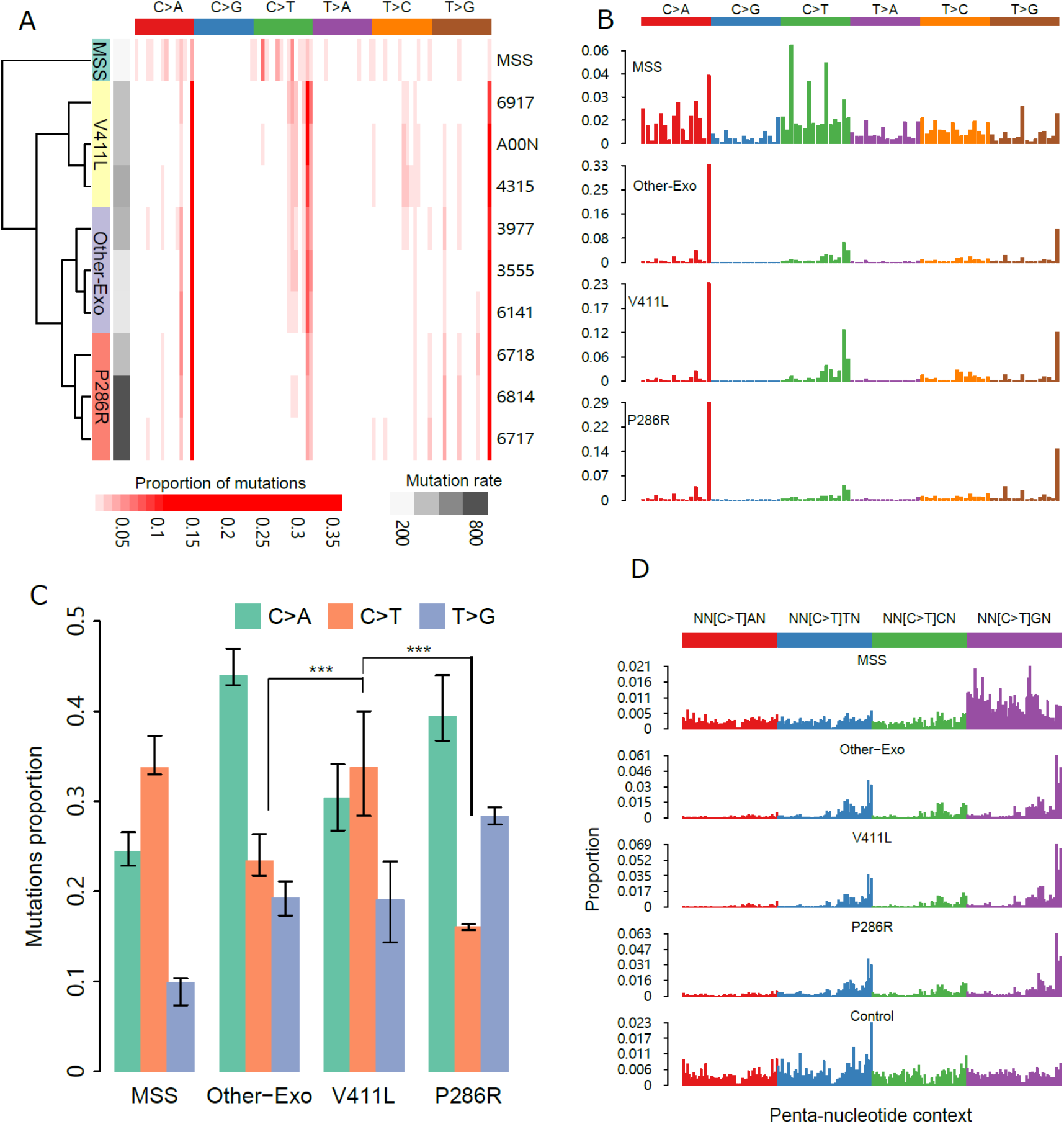
Mutational spectrum of distinct colorectal cancer POLE mutants. **(A)** Hierarchical clustered heatmap of the frequency of 96 types of mutational contexts within each WGS POLE mutant ranging from light red (0%) to dark red (35% of all mutations). Four groups “MSS (microsatellite stable), V411L, P286R and Other-Exo” were labeled on the far left panel, and total mutation burden was indicated in the right bottom. The MSS spectrum is averaged across the 44 TCGA MSS POLE wild-type WGS samples while the TCGA samples ID for each POLE mutant is shown **(B)** Mutational spectrum of four POLE-mutant groups based on 96 mutational contexts, with mutation type indicated on the top panel. **(C)** Proportion of C>A, C>T and T>G mutation in four POLE mutant groups. The significance was calculated by paired Chi-squared test. Error bars represent +/− 2 SE. **(D)** Profile of C>T mutations in penta-nucleotide contexts, with genome-wide frequency of each penta-nucleotide indicated at bottom. *** denotes *P* < 0.001.

We then computed proportions of C>T mutations in different penta-nucleotide contexts to investigate differential enrichment of these mutations (**Figure 1D**). All three types of POLE mutants display enriched C>T mutations in CpG contexts, with the proportion significantly higher in V411L (52.14%), compared with P286R (41.17%) and other POLE mutants (45.67%) (p < 0.0001, Chi-square test, **Figure 1-figure supplement 2**). We also explored the penta-nucleotide context enrichment of C>A and T>G mutations, but did not find substantial differences in the frequency of these mutations in different mutants (**Figure 1-figure supplement 3**). Based this analysis, we can conclude that there are differences in the mutation spectra between the POLE mutants which can be largely attributable to different frequencies of C>T and C>A mutations and the relative frequency of C>T mutations at CpG dinucleotides.

### Methylation and mutation associations in different POLE mutants

Given that we found differences in the frequency of C>T mutations at CpG dinucleotide which is a site subject to DNA methylation we sought to compare the relationship between 5-methylcytosine (5mC) level and C>T mutation frequency in the different POLE mutants. A significant association between methylation and mutation burden has been shown in POLE mutants, with more mutations in highly methylated sites, indicating the presence of 5mC as a contributing factor in POLE-associated mutagenesis (Poulos et al., 2017). In this study, we first investigated methylation levels at CpG dinucleotides by using whole genome bisulfite sequencing data of normal sigmoid colon, and correlated the methylation level with mutational burden across the colorectal cancer genome. In all three types of POLE mutants, including POLE wild-type MSS samples, the mutation burden increased significantly with methylation levels (**Figure 2A**). To investigate whether differences in the CpG mutation load between the different POLE mutants is dependent on methylation level, mutation burden within each bin of methylation level for the different POLE mutants were normalized against V411L. We found that the slope of the normalized mutation burden does not substantially deviate from zero across increasing levels of methylation (**Figure 2B**). This finding suggests that, while mutation burden at CpG sites are dependent on cytosine methylation levels, there is also a cytosine methylation-independent component of mutation load that accounts for the relatively higher number of C>T mutations in V411L compared with the other POLE mutants.

**Figure 2.**
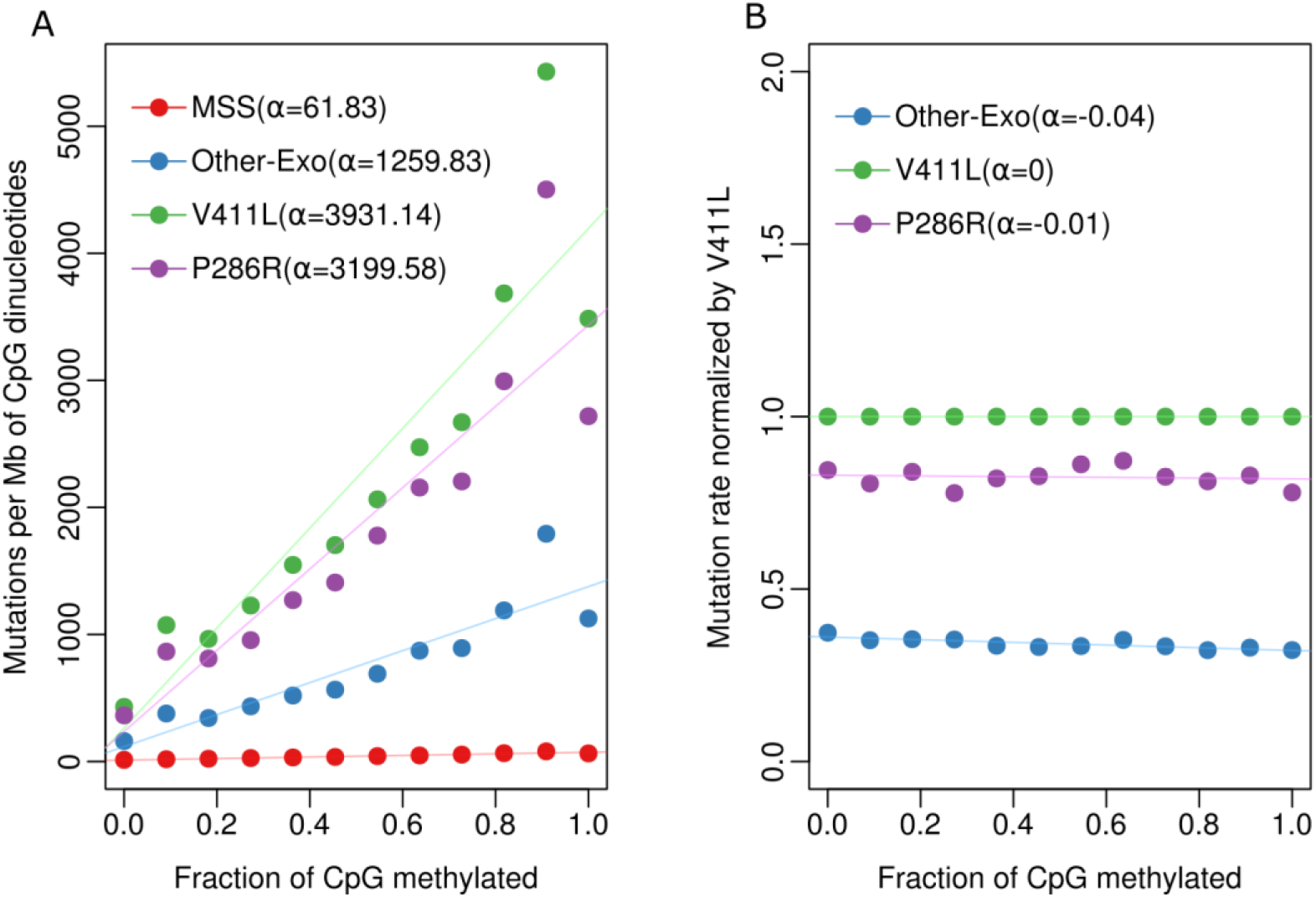
Association of methylation and mutation in different POLE mutants. **(A)** Correlation between mutations per megabase (Mb) at CpG dinucleotides and fractions of CpGs methylated across different POLE mutants and microsatellite stable (MSS) samples. **(B)** Mutation burden of each mutant was normalized by the mutation rate of V411L in each methylation level.

Finally, in all POLE mutants, mutation burden peaks when the level of CpG methylation measured is between 90% and 100%, but decreases when the level of CpG methylation level is equal 100%. We examined whether sequencing coverage, replication timing or repeat sequences in different methylation levels contributes to this change, but found that they were not associated with this observation (**Figure 2-figure supplement 1**). We then tested the composition of penta-nucleotide contexts at different levels of methylation, since C>T mutations are also enriched in specific penta-nucleotide context as discussed above (**Figure 1-figure supplement 2**). We found that there are more CpGs in the TTCGN context in the 90-100% CpG methylation bin compared with the 100% CpG methylation bin, accounting for 8.67% and 5.62% of CpGs respectively (p<0.001, Chi-square test, **Figure 2-figure supplement 2**). Following normalization for penta-context composition across the different bins, the mutation rate at the 90-100% bin decreased by 17.6% (Other-Exo), 18.04% (V411L) and 20.02% (P286R) POLE mutants respectively, making the mutation rate in this bin more similar to that of the 100% methylated CpG sites (**Figure 2-figure supplement 3**). This finding again demonstrates that different preferences for penta-nucleotide context within POLE mutants can account for differences in the observed mutational patterns.

### Distinct POLE mutants show similar genome-wide mutational patterns

#### Characterization of mutations in CTCF binding sites

CCCTC-binding factor (CTCF) is a transcription factor and plays an essential role in constructing three-dimensional genome organization. Somatic mutations in CTCF binding sites of the CTCF-cohesin complex (CBS) are widely observed in cancer genomes (Kaiser et al., 2016). Samples with *POLE* mutations displayed lower mutation frequencies at, and adjacent to, CBS when compared with flanking regions (Katainen et al., 2015), but the mutation rate of distinct POLE mutants has not been examined. We calculated mutation counts at each position within 1000 nucleotides from the CTCF motif center and we identified a distinct pattern whereby mutation burden was significantly decreased in all the three mutants (**Figure 3A**). For each mutant, mutation load starts to decline approximately 110 nucleotides from the CTCF motif center, and then presents a significant lower mutation frequency than expected by chance within the center of the CTCF motif, especially at the central cytosine nucleotide (P<0.001, paired Wilcoxon signed-rank test, **Figure 3-figure supplement 1A, B**). We characterized the mutation signature within this ±110 nucleotide region and we observed a similar mutational pattern with genome-wide signature (**Figure 3-figure supplement 1C**), suggesting that at least some of the CBS sites examined are still under the POLE mutation process.

**Figure 3.**
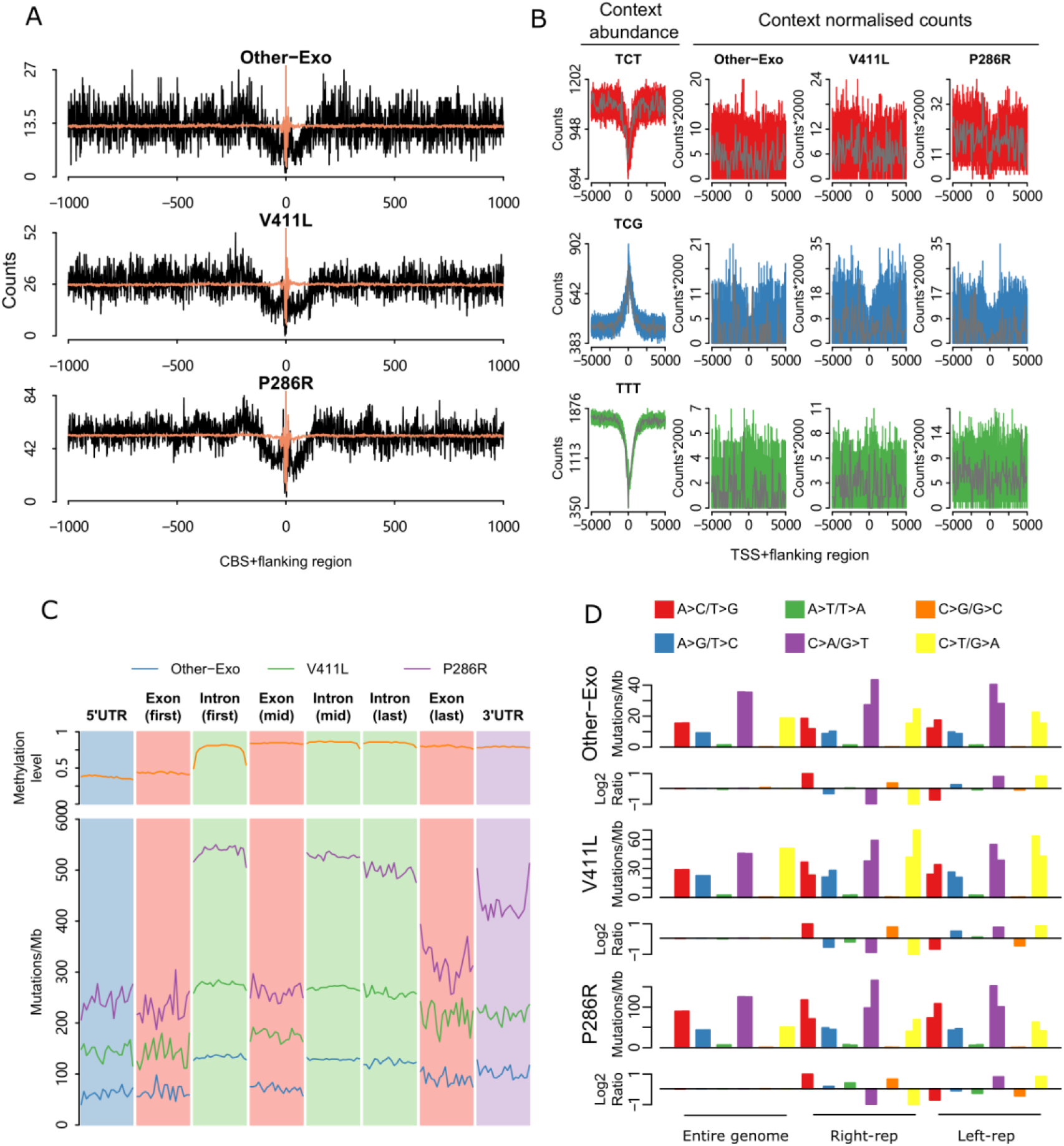
Genome-wide mutational patterns of distinct POLE mutants. **(A)** Somatic substitutions at CBSs with a flanking sequence of 1 kilo bp in different POLE mutants. The expected mutation was indicated in light red colour. **(B)** Mutation profile around transcription start sites in different mutants. Three primary mutation types C>A (red), C>T (blue) and T>G (green) in specific context were showed. Mutation counts were normalized by the number of corresponding context and the abundance of each context was displayed in the far left panel, together with mutation data in 100 bp bins (grey) is shown. **(C)** Profile of mutation burden across different part of genes in different mutants. Each part of gene was divided into 20 bins and mutation burden was calculated separately. Methylation level of each part was showed in the top panel. **(D)** Mutational strand asymmetry associated with replication in different mutants. Lower panel of each mutant shows the log2 ratio of each pair of bars.

#### Mutation density around the transcription start site

We also investigated mutation density around the transcription start site (TSS) in different POLE mutants. The DNA sequence around the TSS can show distinct mutation patterns, as active promoters are occupied with transcription factors, which may inhibit DNA repair access or activity (Perera et al., 2016, Sabarinathan et al., 2016). We examined mutation profiles of C>T, C>A and T>G mutations around TSSs for each POLE mutant. Notably, before normalization, T>G mutations were substantially decreased at the TSS (**Figure 3-figure supplement 2**). However, following normalization for trinucleotide sequence context, this was no longer evident, and we only observed substantial decrease in C>T mutations close to the TSS likely due to reduced DNA methylation at many gene promoters (**Figure 3B**)

#### Exonic regions show decreased mutation burden in POLE mutants

Increased mismatch repair (MMR) activity at exons compared with introns has been shown to result in a significant decrease in exonic mutation rate in MMR proficient POLE mutants (Frigola et al., 2017). We investigated mutation patterns of exonic and intronic regions in different POLE mutants (**Figure 3C**). All three kinds of POLE mutants showed decreased mutation rates in exonic region. Particularly in P286R mutants, the average mutation burden in the middle of intronic regions is approximately double the count in the middle of exonic regions (260 vs 528 Mut/Mb, **Figure 3-figure supplement 3**).

#### POLE mutants present mutational strand asymmetries

Since the exonuclease domain of POLE is responsible for proofreading during synthesis of the DNA leading strand, mutations caused by deficiency of the domain should show very strong strand asymmetries (Haradhvala et al., 2016). We identified this phenomenon in all distinct POLE mutants, with all mutants showing similar levels of strand asymmetry (**Figure 3D**). As expected, in left (5’)-replicating regions that are enriched in leading strand synthesis we observed C>A, C>T and T>G mutations predominantly. On the contrary, G>T, G>A and A>C mutations are predominantly in right (3’)-replicating regions that are enriched in lagging strand synthesis.

#### Periodicity of mutation rate across and within nucleosomes

The minor groove of DNA that wraps around nucleosomes presents a differential pattern due to its physical interaction with histones, and this pattern determines periodicity in mutation rate (Brown et al., 2018, Pich et al., 2018). Colorectal cancers with contribution from signature 10 have been reported to exhibit a positive minor-in relative increase of mutation rate as a consequence of interaction between the processes of DNA damage and repair within the nucleosome (Pich et al., 2018). We investigated mutation rate periodicity in each specific POLE mutant separately, and we observed the positive minor-in relative increase of mutation rate in all POLE mutants to comparable levels, suggesting that the different POLE mutant induced mutations are not differentially affected by DNA-histone interactions (**Figure 3-figure supplement 4**).

#### Increased mutation rate at late replication timing

Finally, the mutation rate in late-replicating regions should be higher than in early-replicating regions in MMR proficient cancer samples due to differential MMR efficiency (Supek and Lehner, 2015). Although all POLE mutants showed high mutational burden, they are MMR proficient with a microsatellite stable phenotype. The mutation burden of a range of mutational signatures have been associated with DNA replication timing, and a significant correlation with replication timing has been reported in cancer samples with POLE mutant associated mutational signature 10 (Tomkova et al., 2018). We calculated mutation rate in genomic region with distinct replication timings. As expected, all mutants similarly displayed higher mutation rates in late-replicating regions than in early-replicating regions despite their different mutational context (**Figure 3-figure supplement 5**).

### Mutational context of POLE-mutants predisposes colorectal cancers to developing TP53 R213* mutation hotspots

Since we had identified that different POLE mutants have different mutation spectra, we next sought to determine whether this may predispose cells to specific additional cancer driver mutations. We screened a list of 47 cancer driver mutation hotspots, determined based on recurrence in cohorts where we could access mutation calls in an unbiased manner (see **Methods**), in a total of 7,345 colorectal cancer samples including 47 POLE mutants (16 P286R, 15 V411L and 16 Other-Exo mutants with Sig10) and 7,298 POLE wild-type samples, to investigate if any hotspots are enriched in specific POLE mutants (**Supplementary table 1**). For all hotspots tested, only the truncating mutation R213* in TP53 was identified to be significantly enriched in POLE P286R mutants (P= 0.0076, Fisher’s exact test, Benjamini-Hochberg FDR 10%, **Figure 4A, Supplementary table 2**). For all P286R mutants, 62.5% (10/16) harbor this hotspot, while it occurs in only 19.4% (6/31) of other POLE mutants (**Figure 4A, Supplementary table 3**). For the remaining 7,298 POLE wild-type samples, only 2.2% (163/7298) were identified with this hotspot mutation. This nonsense mutation is a C>T transition in the context of TT[C>T]GA (**Figure 4B**), which is a relatively enriched context in P286R mutants, with adenine being more prevalent in the 5^th^ position, compared with the other POLE mutants (**Figure 4C**, p <0.05, Student’s t-test, **Figure 4-figure supplement 1**). As the relative frequency of the TT[C>T]GA pentanucleotide context differs between individual samples, we compared its relative frequency in POLE mutants with and without the TP53 R213* mutation. This mutational context was found to not only be significantly higher in POLE P286R mutants with the TP53 R213* mutation compared with those without this mutation (P = 0.0088, Student’s t-test, **Figure 4D**), but also significantly higher across all POLE mutants with the TP53 R213* mutation compared with those without this mutation (P = 0.0161, Student’s t-test, **Figure 4E**). This suggests that for C>T mutations, POLE mutant colorectal cancers (and in particular the P286R mutant) that generate relatively more mutations in the TT[C>T]GA context, have a higher chance of acquiring the TP53 R213* mutation.

**Figure 4.**
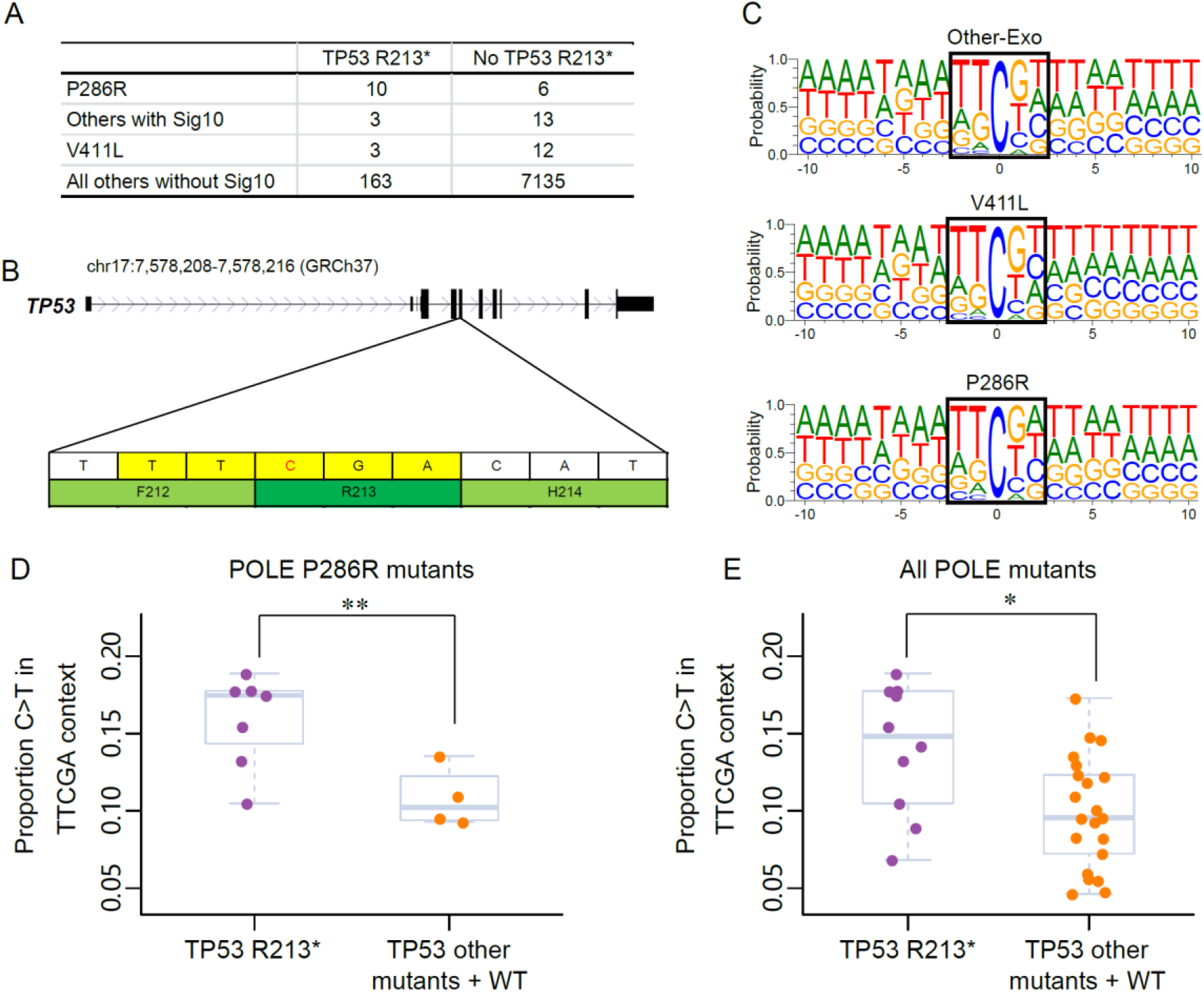
Mutation hotspots in POLE mutants. **(A)** Contingency table of different POLE-mutant and POLE wild type colorectal cancer samples with or without the TP53 R213* mutation. Samples with Sig10 were confirmed by either POLE driver mutation or mutational spectrum clustering. **(B)** Truncating mutation TP53 R213* was caused by C>T substitution in TT[C>T]GA context. **(C)** Frequency of 21-bp sequence context centered by mutated cytosine in different POLE mutants, and the penta-nucleotide contexts were indicated in black box. Proportion of C>T mutations in the TTCGA pentanucleotide context in POLE P286R mutants (D) and all POLE mutants **(E)** with and without the TP53 R213* mutation. * < 0.05. ** < 0.01, Student’s t-test.

## Discussion

In this study, we investigated genome-wide regional mutational profiles of different POLE mutants, as well as their influence on driver mutation formation in cancer. Genomes with POLE functional defects present with differential mutation spectra but show largely similar regional mutational profiles. Significantly, we identified a recurrent nonsense mutation in TP53 that is enriched in P286R mutants, indicating a new insight into mutational processes of specific POLE mutants.

All POLE mutants had frequent C>A, C>T and T>G mutations in TCT, TCG and TTT contexts respectively, which is consistent with the classic POLE signature. However, we found that POLE V411L mutants carry significantly more C>T mutations genome-wide than other mutants (V411L vs P286R, P<0.001, Chi-squared test). V411L and P286R are the two most frequent POLE mutants and they are located far away from each other in the exonuclease domain, thus conferring different functions (Palles et al., 2013). P286 lies in the DNA binding pocket, which might interact with single strand DNA by directly perturbing the binding pocket. However, V411 is far away from exonuclease active site may function by affecting the positions of other residues adjacent to the active site (Briggs and Tomlinson, 2013). In yeast, POLE mutants with a weak exonuclease activity have more C>T and less C>A mutations than mutants with no exonuclease activity (Xing et al., 2019). We therefore speculate that the proportionally reduced C>A and increased C>T mutation loads in V411 may arise due to stronger exonuclease activity, as the mutation is distal from that site. Consistent with this, in a cell free system, V411L was found to have 3-fold reduced exonuclease activity compared to wild-type, while P286R mutants displayed a 10-fold reduction (Shinbrot et al., 2014).

Methylated cytosine have been shown to readily mutate to thymine as a result of methylcytosine deamination (Sassa et al., 2016). We found that although V411L had comparably more mutations at the cytosine of CpG dinucleotides than P286R and other POLE mutants (16.68% (V411L), 6.38% (P286R) and 10.19% (Other-Exo)), all mutants showed the same positive association with methylation level after adjusting for total CpG mutation count. Therefore, although methylation is an important determinant of CpG mutagenesis, the CpG sequence context is particularly favorable for mutagenesis in the V411L mutants. In addition, we observed unusually high mutation rate at methylation levels between 90-100%. As most C>T mutations are enriched in the specific TTCGN context, we demonstrate that the composition of contexts in different methylation level can influence the mutation burden.

Altered *POLE* enzyme function could result in substantial mutation accumulation, and we observed uneven distribution of mutations at local regions in different mutants. CBSs are frequently mutated across different cancer types, and are a major mutational hotspot in noncoding cancer genomes (Katainen et al., 2015). CTCF binding sites display a specific mutation pattern in skin cancers due to differential nucleotide excision repair (Poulos et al., 2016). We observed decreased mutation density at and adjacent to CBSs, and the decline starts at around 110 nucleotide distance from the center of the CBS. It has been proposed that the decrease in mutation density in this region might be due to either the use of an alternative polymerase (Katainen et al., 2015). CTCF-cohesin binding sites might be treated like DNA-protein crosslinks during replication, so they get bypassed with the help of an accessory helicase RTEL1 and is later filled by translesion synthesis (Sparks et al., 2019). Finally, disparity in mutation rate between exon and intron regions, and early and late replication timing regions were identified in all POLE mutants, although the effect appeared strongest in the P286R mutants, possibly due to the higher mutation burden in these samples. These results suggest that mismatch repair is an important system to protect against POLE replication errors regardless of subtle differences in the way the mutations were generated. Overall, in-depth analysis of mutation patterns with respect to a vast array of genomic and epigenetic features revealed largely unchanged mutation patterns between different POLE mutants suggesting that these factors that affect mutagenesis do not strongly interact with the biochemistry of the enzyme to modify their mutagenic effects.

Mutational signatures representing the spectrum of different somatic mutations can be employed to decipher the mutational process that operated within an individual cancer (Alexandrov et al., 2013b). Recent studies have revealed the associations between mutational processes and somatic driver mutations to some extent, and indicated that altered tri-nucleotide preferences arising from a certain signature would increase the likelihood of the associated driver mutation arising (Poulos et al., 2018, Temko et al., 2018). Previous studies have identified an association between the TP53 R213* truncating mutation and POLE mutant cancers (Shinbrot et al., 2014, Poulos et al., 2017). Our study has further identified that this TP53 hotspot is significantly enriched in POLE P286R mutants (62.5%) in colorectal cancer. The TP53 R213* truncating mutation is caused by a C>T transition in a TTCGA penta-nucleotide context, and we found that POLE mutants with this mutation do generally have higher relative frequency of this mutational context compared to POLE mutants without this mutation. This implies a possible direct causal relationship between POLE-associated mutagenesis and acquisition of this driver mutation.

In summary, understanding how specific driver mutation may arise could lead to new targeted therapeutic strategies. This study has shown the importance of further subtyping cancers, not only focusing on the mutated genes, but also the specific mutations within those particular genes. Stratifying samples based on DNA polymerase activity defects has enabled us to gain a better understand the mutational processes in colorectal cancer genomes.Materials and methods Somatic mutations and sample classification

## Materials and methods

### Somatic mutations and sample classification

All somatic mutations of 53 whole genomes colorectal cancer were obtained from The Cancer Genome Atlas (TCGA) (Wilks et al., 2014). Microsatellite status and *POLE* mutation status were provided for each sample as listed in **Supplementary table 4**.

2,506 colorectal samples with complete whole exome/target capture data from TCGA and previously published datasets (Giannakis et al., 2016, Seshagiri et al., 2012, Yaeger et al., 2018) were first used to identify recurrent driver mutation sites (in at least 20 individuals) in colorectal cancer. Furthermore, 257 whole genome sequenced colorectal cancer samples but with only selected mutation data available (Katainen et al., 2015) and another 4,582 colorectal samples also with target capture data from AACR Project GENIE through cBioPortal (Cerami et al., 2012) were additionally used for analyzing POLE mutants with driver mutation hotspots. A table showing the sample cohorts and the mutation status of all samples are show in **Supplementary tables 5 and 1**, respectively. Mutations were annotated by oncotator-1.9.9.0 when necessary.

For all samples with non-silent mutations in *POLE*, we performed clustering based on proportion of 96 tri-nucleotide context in order to distinguish functional POLE mutants that are characterized by mutational signature 10. For samples obtained from AACR Project GENIE, functional POLE mutants were confirmed by a list of reported driver mutations reported previously (Campbell et al., 2017).

### Mutational signature analysis

The profile of each signature was displayed using the six substitution subtypes: C>A, C>G, C>T, T>A, T>C and T>G. For signature generated by tri-nucleotide context, each substitution was examined by incorporating information on the bases immediately 5’ and 3’ to each mutated base to generate 96 possible mutation types. For signature generated by penta-nucleotide context, each substitution was examined by incorporating information of two nucleotides at 5’ and 3’ to each mutated base resulting in 1536 possible mutation types. The mutational signatures were displayed and reported based on the observed tri-nucleotide/penta-nucleotide frequency of the human genome.

### Methylation data and mutation

Whole genome bisulfite sequencing data from normal sigmoid colon tissue were downloaded from the Roadmap Epigenomics Atlas (Kundaje et al., 2015). All cytosines in the CpG di-nucleotide were merged into 12 bins according to their methylation level as: [0], (0, 0.1], .., (0.9, 1.0), [1]. These bins were then used as intersected regions to calculate mutation rate in each methylation level.

### Penta-nucleotide context normalization for each methylated level

First, the abundance of each penta-nucleotide context in which CpG context located were calculated by using the downloaded whole genome bisulfite sequencing data. Then we weighted each context (*f*) by their counts, and made the sum of weights values equal to 1. Similarly, the abundance of penta-nucleotide context in each methylation level was also calculated and weighted (*F*). Next, the counts of mutated contexts of each sample were computed (*N*). Finally, the normalized value of each methylation level was obtained as (*C*):

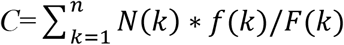

### CTCF motifs and data analysis

CTCF/cohesin binding sites for the LoVo cell line were obtained from published paper(Katainen et al., 2015). Each CTCF motif was extended to 1000 bp on each side, and the mutation profiles were generated by counting mutations that are intersected with these sequences at each base. In order to obtain expected counts that are affected by fraction of distinct contexts, the following procedures were conducted. First, the count (*M*) of each mutated context was calculated in the overall extended sequences. Then, the abundance of each context in the whole extended sequence was computed as A, and for each base of each line in the stacked sequence, the relative frequency (*f*) was calculated by dividing the number of mutations with that context by the abundance *A*.

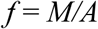

Next, for each position *p* within each sequence, we weighted *p* by its respective context-specific frequency *f*, and made the sum of weights across all 2001 values equal to 1. So the vector of weights *W_p_* across the specific 2001-bp sequence is given by:

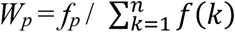

Subsequently, all 2001-bp sequences were stacked and the expected count at position *p* was computed as *m*W_p_*, where *m* is the count of mutations in the sequence where *p* is located. Finally, the expected count at a given position *p* of the stack of aligned sequences is obtained as the sum of all the expected counts at each sequence of position *p*.

### Generation of mutation and profiles across transcription start sites

The information of transcription start sites (TSS) were obtained from canonical genes from the UCSC table browser. For each set of TSSs, mutation profiles were generated by counting the number of three major types of mutations (C>A, C>T and T>G) across a ±5,000-bp window centered by TSS. Mutation counts were normalized by dividing the abundance of corresponding context at each position.

### Periodicity of the relative increase of mutation rate

The methods used for mutational periodicity analysis referred to previously published paper and script (Pich et al., 2018). Briefly, 147 bp length mid-fragments of high-coverage MNase-seq reads representing putative nucleosome dyads were obtained from the paper published by Gaffney et al (Gaffney et al., 2012). Then the wig format file was converted to the bed format for following analysis. The relative positions to the dyad of two center nucleotides of the DNA to decide the minor groove facing the histones and away were obtained from the paper published by Cui and Zhurkin (Cui and Zhurkin, 2010). These positions combined with somatic mutation data were used to calculate mutation rate in stretches of DNA with the minor groove facing histones and away from them.

### Calculating mutational strand asymmetries

Replication direction was defined using replication timing profiles that are from previously published paper (Koren et al., 2012). Left- and right-replicating regions were determined by the derivative of the profile, assigning negative and positive as left-replicating and right-replicating respectively. For a given mutation type in specific replication direction, the mutation counts (*N*) in that region were calculated, and its complementary mutation was calculated as *n*. Then, asymmetry (*A*) was calculated in a given region by:

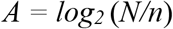

### Computing mutation density in exonic and intronic region

All gene coordinates were obtained from UCSC table browser. Each gene was divided into eight parts as 5’UTR, first exon, first intron, middle exon, middle intron, last intron, last exon and 3’UTR. As the length of sequence in each part is various, we divided every sequence into 20 bins in a given part. Sequences with length of less 20-bp were discarded. Then, the mutation density was calculated and normalized to mutations per Mb and plotted in each bin.

### Replication timing and mutation rate

The replication time of different chromosome position was obtained for the HepG2 cell line from the ENCODE data portal (Sloan et al., 2016). All the sequence with known replication time was integrated into 5 bins from late to early: [-4.51712, 30.8225), [30.8225, 44.19), [44.19, 55.8262), [55.8262, 63.7717), [63.7717, 80.6964]. Mutations located in each bin were calculated.

## Author Contributions

H.F. carried out overall analysis of data. H.F., J.A.B. and R.C.P. were involved in experimental design. R.K and L.A.A. contributed data analyses and datasets. J.W.H.W undertook overall project design and conceptualized the study. H.F. and J.W.H.W wrote the manuscript with contributions from the other co-authors.

---

**Figure 1-figure supplement 1.**
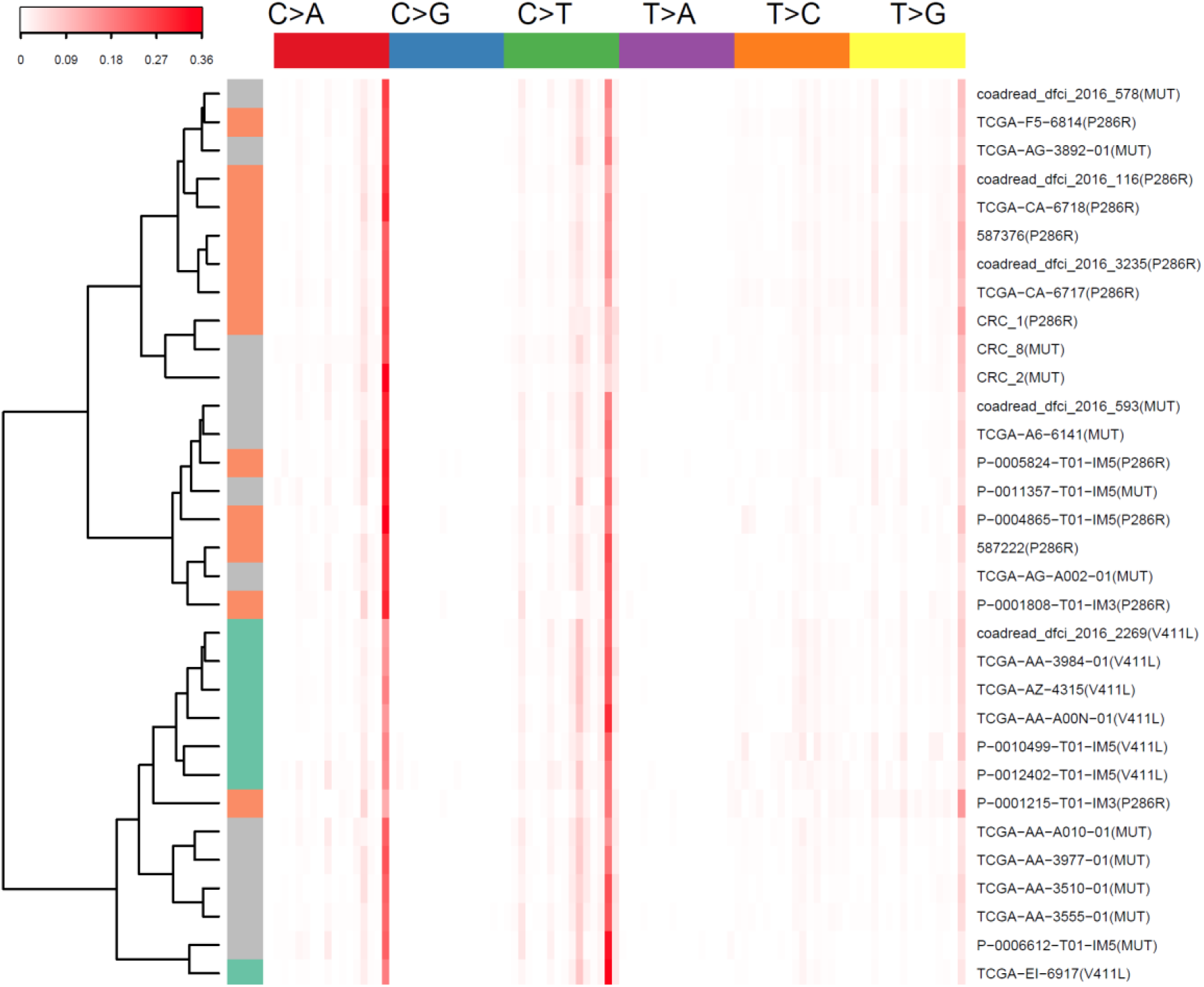
Hierarchical clustered heatmap of the frequency of 96 types of mutational contexts for 32 POLE samples that have been whole genome, whole exome or targeted sequenced.

**Figure 1-figure supplement 2.**
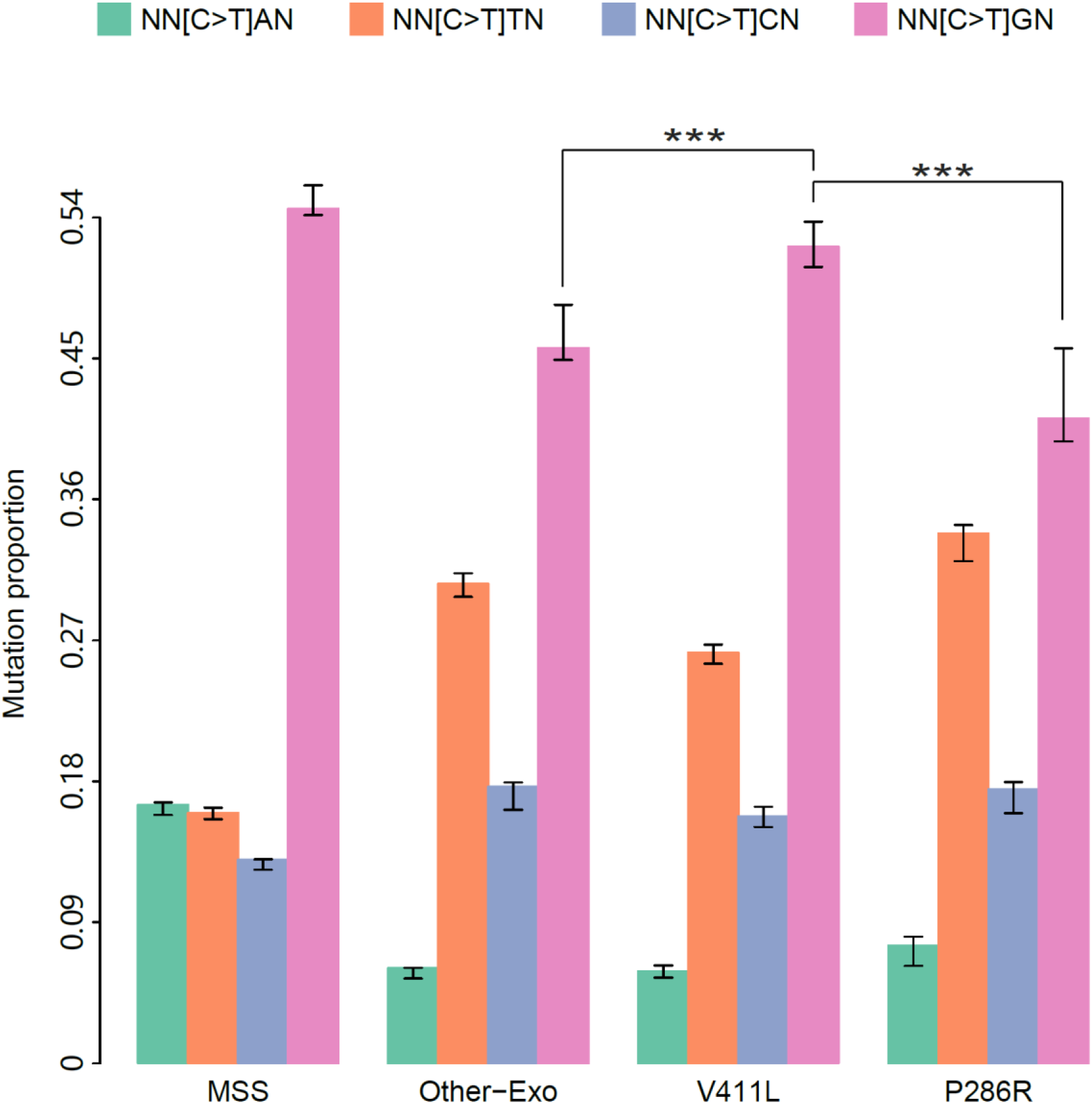
Proportion of C>T mutations in the CpA, CpC, CpG and CpT contexts.

**Figure 1-figure supplement 3.**
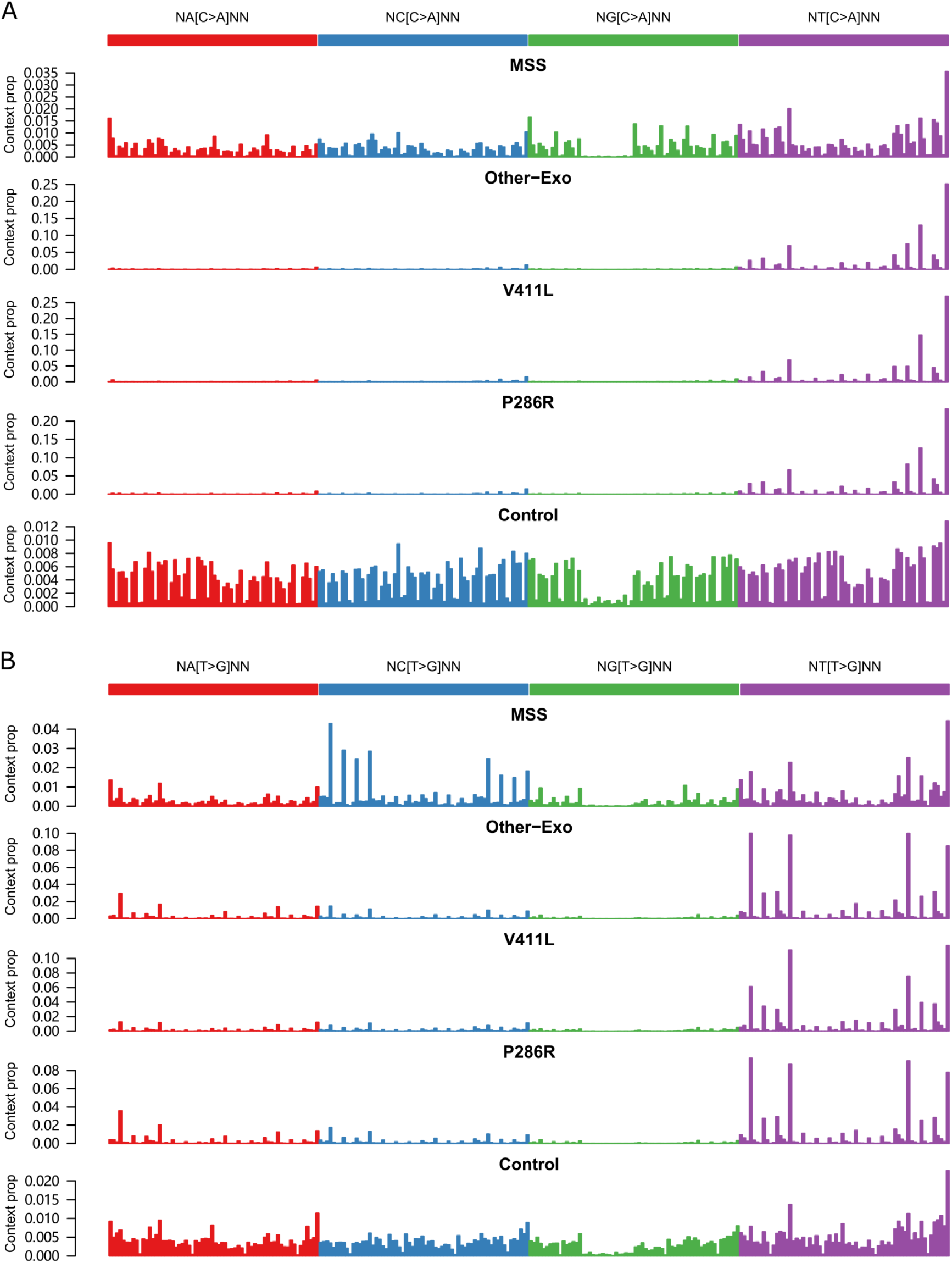
Profile of (A) C>A and (B) T>G mutations in penta-nucleotide contexts, with genome-wide frequency of each penta-nucleotide indicated at bottom of each figure.

**Figure 2-figure supplement 1.**
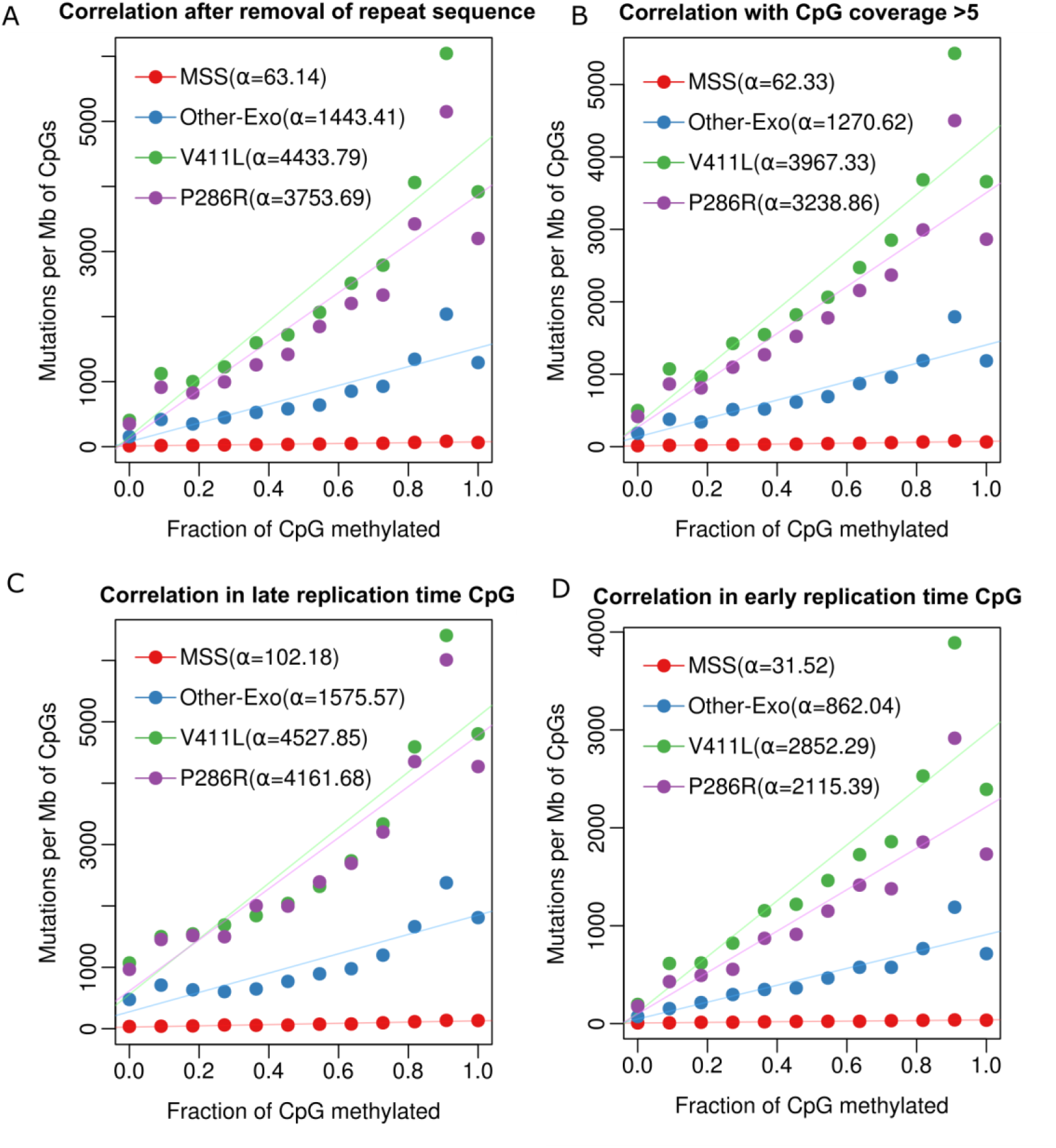
Association of methylation and mutation in different POLE mutants in different condition. **(A)** Correlation of methylation and mutation burden after removal of repeat sequence in CpGs. **(B)** Correlation of methylation and mutation burden in the condition of the coverage of CpGs greater than five. **(C)** Correlation of methylation and mutation burden in late replication timing CpGs. **(D)** Correlation of methylation and mutation in early replication timing CpGs.

**Figure 2-figure supplement 2.**
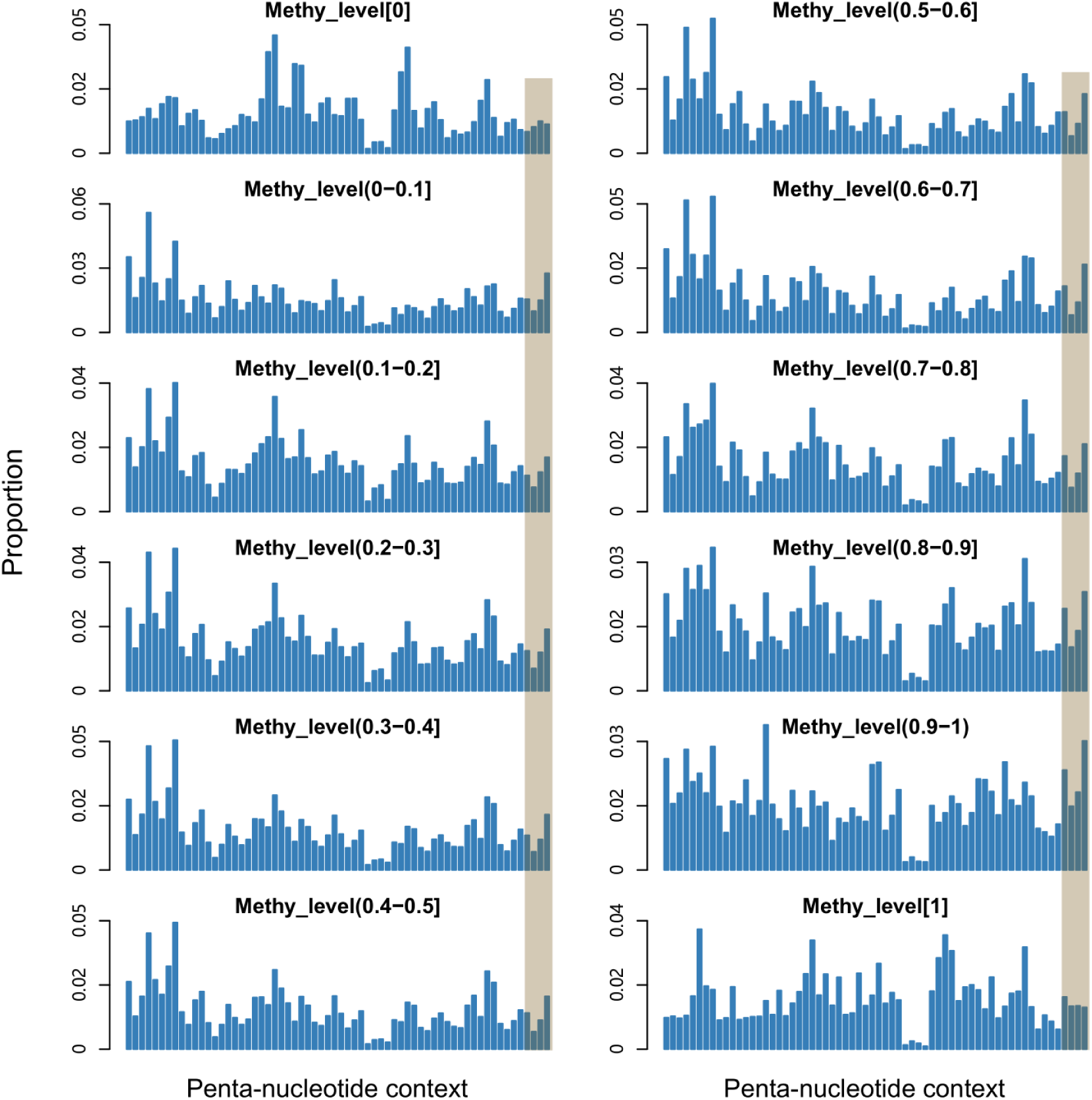
Proportion of each “NNCGN” penta-nucleotide context in different methylation level, with “TTCGN” shadowed.

**Figure 2-figure supplement 3.**
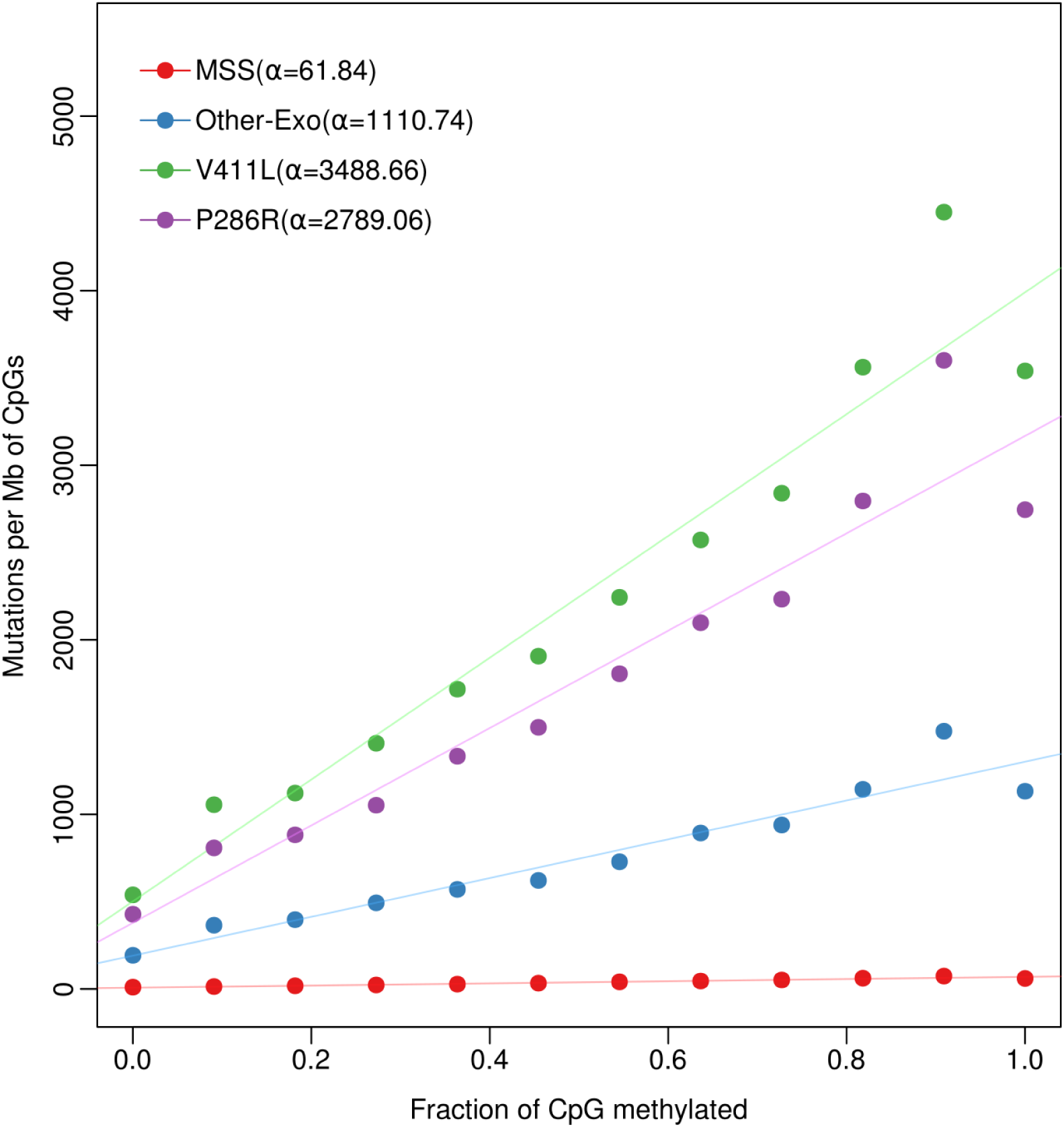
Correlation of methylation and mutation burden after normalization of penta-nucleotide context composition.

**Figure 3-figure supplement 1.**
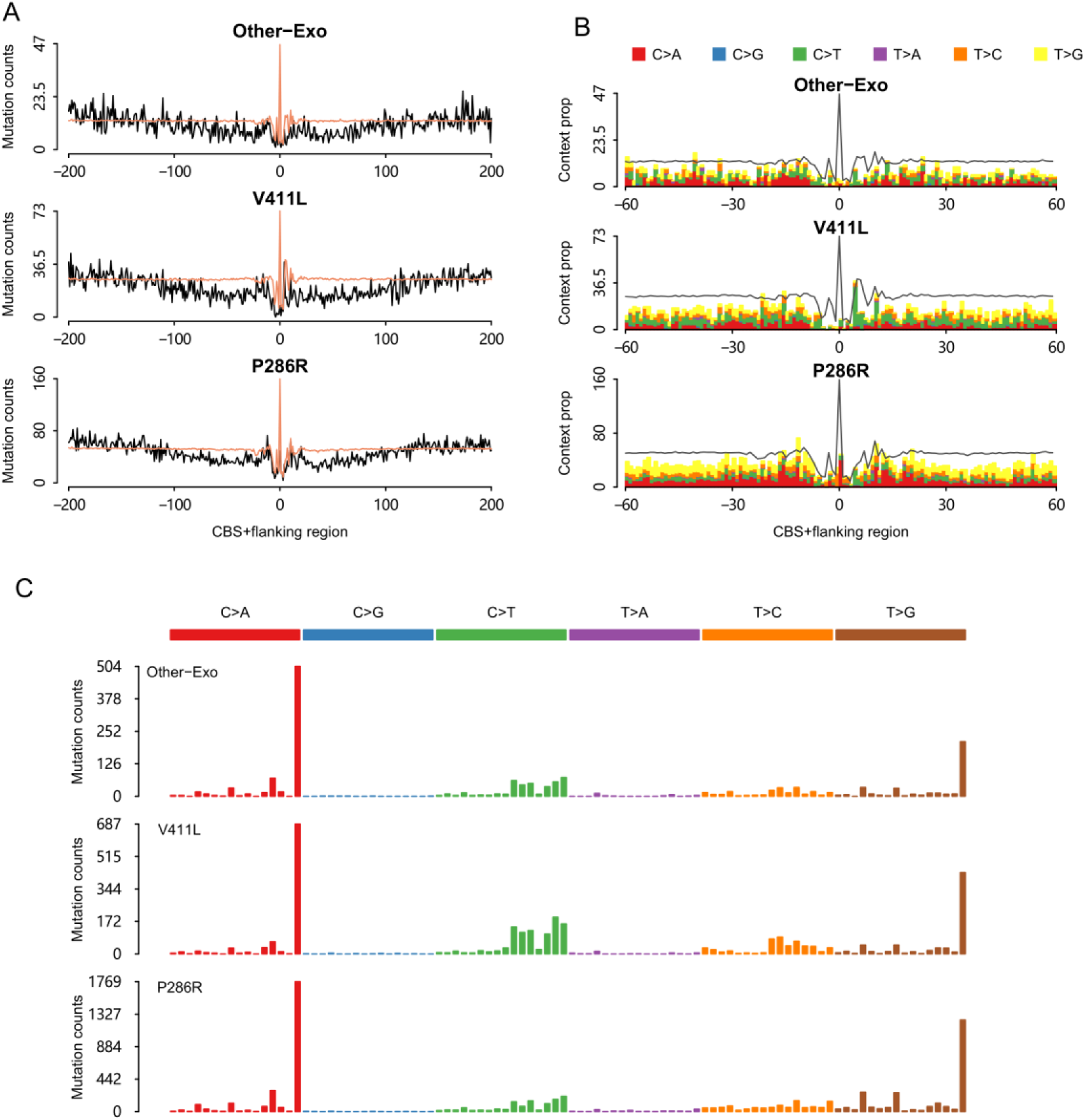
Mutational patterns in and adjacent to CBS region. **(A)** Somatic substitutions at CBSs with a flanking sequence of 200 bp in different POLE mutants. The expected mutation was indicated in light red color. **(B)** Profile of mutation type was showed in CBSs with a flanking 200 bp sequence. **(C)** Mutational spectrum within ± 200bp sequence centered by CBS based on 96 mutational contexts.

**Figure 3-figure supplement 2.**
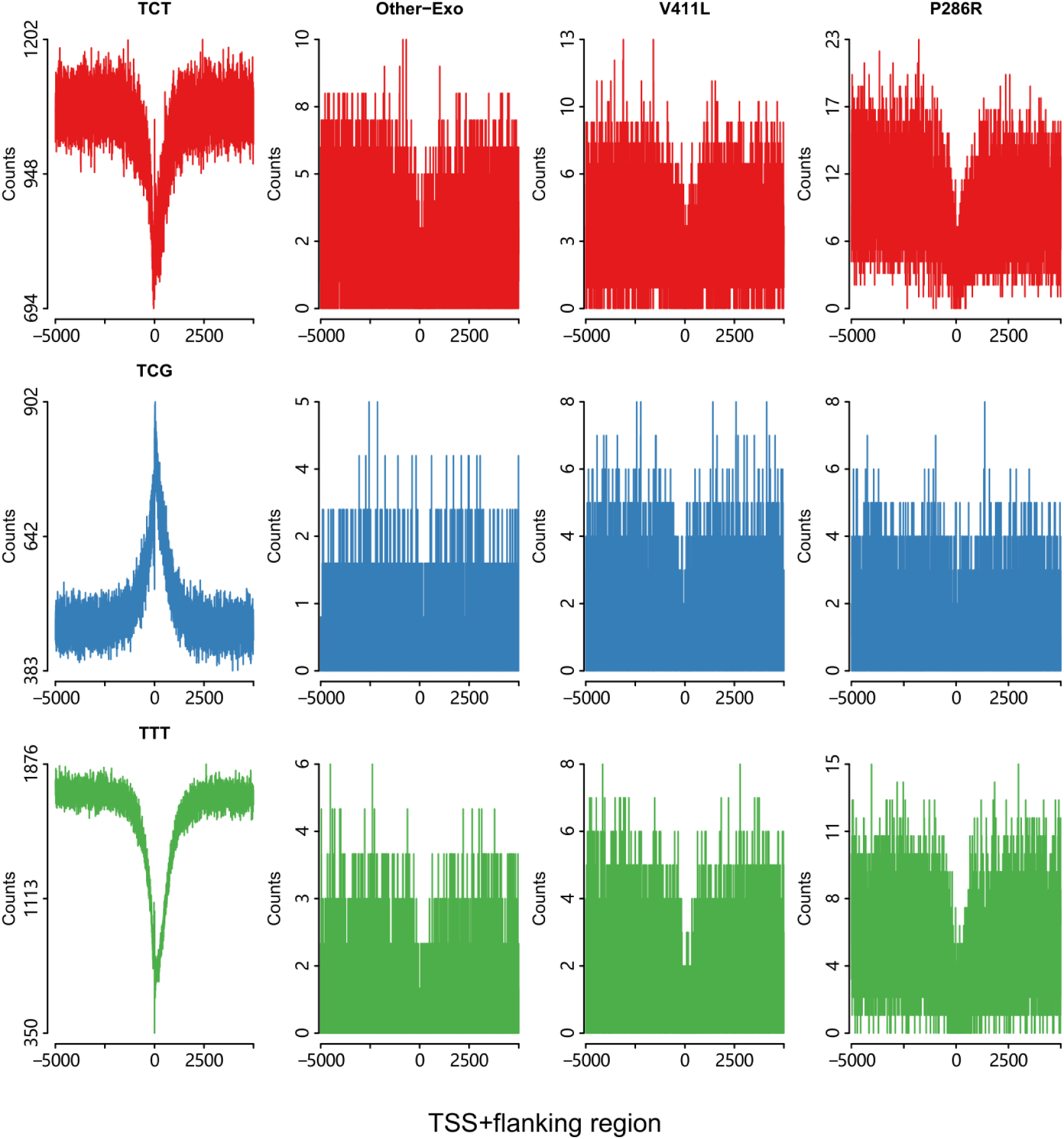
Mutation profile around transcription start sites in different mutants. Three primary mutation types C>A, C>T and T>G in specific context were showed, and the abundance of each context was displayed in far left panel.

**Figure 3-figure supplement 3.**
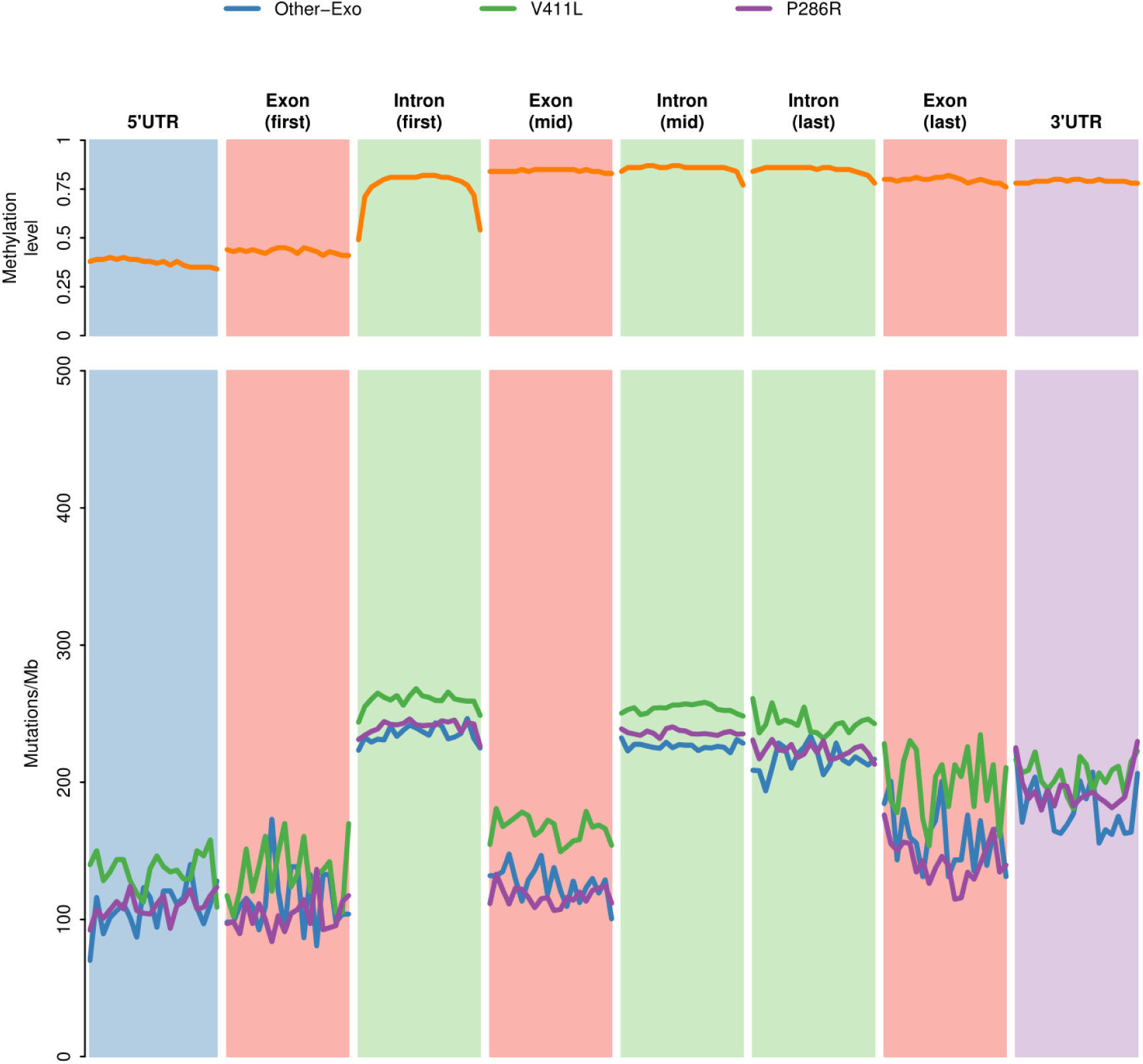
Profile of mutation burden across different parts of genes in different mutants. Mutation burden was normalized by the total number of mutations in each type of mutant.

**Figure 3-figure supplement 4.**
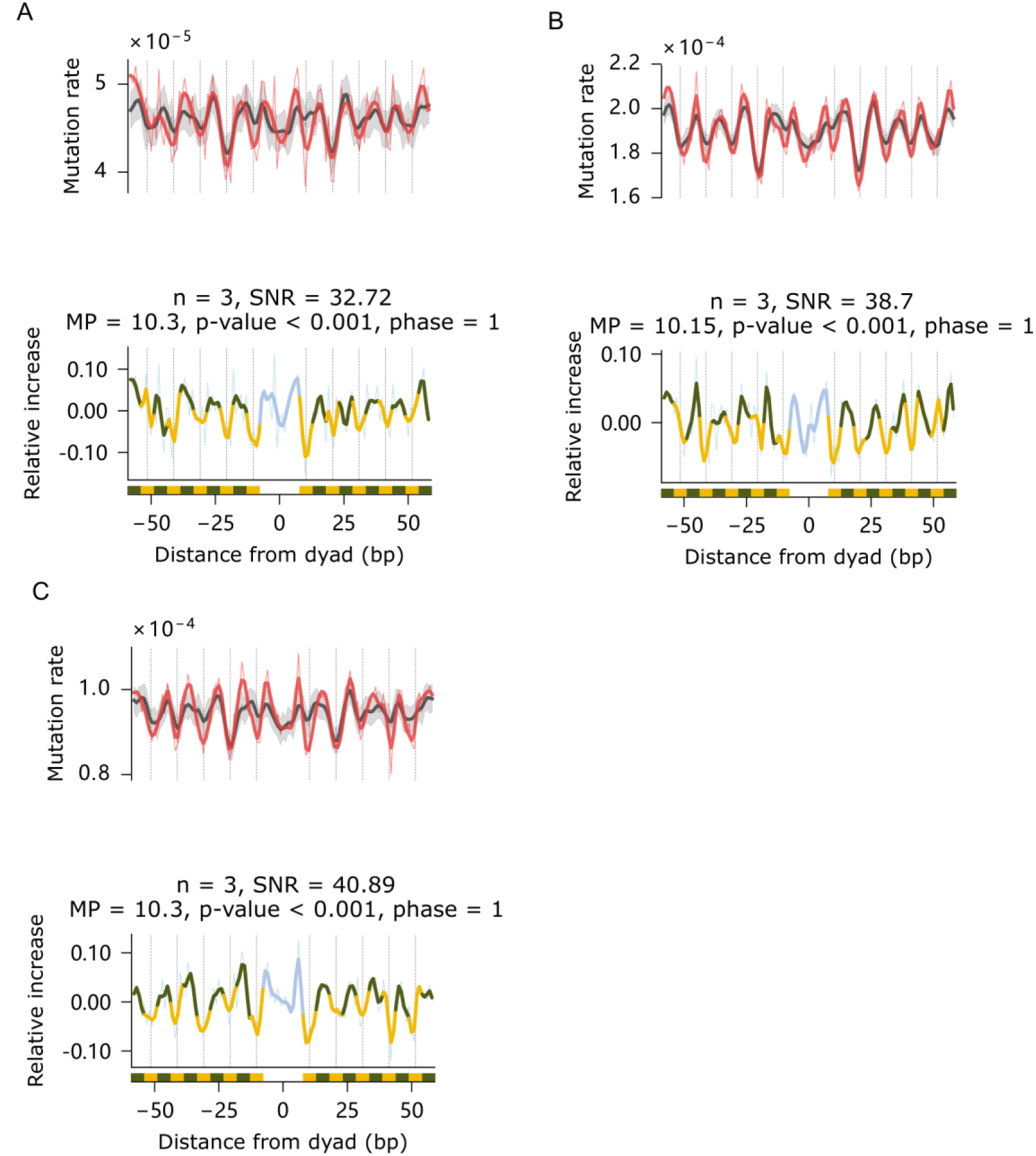
Periodicity of tumor mutation rate within nucleosomes in different mutants: **(A)** Other-Exo, **(B)** V411L and **(C)** P286R. For each figure, the top panel shows observed and expected mutation rate, and the bottom panel shows relative increase of mutation rate. The bottom bar is schematic representation of alternating sequences of DNA with minor groove facing toward and away from histones.

**Figure 3-figure supplement 5.**
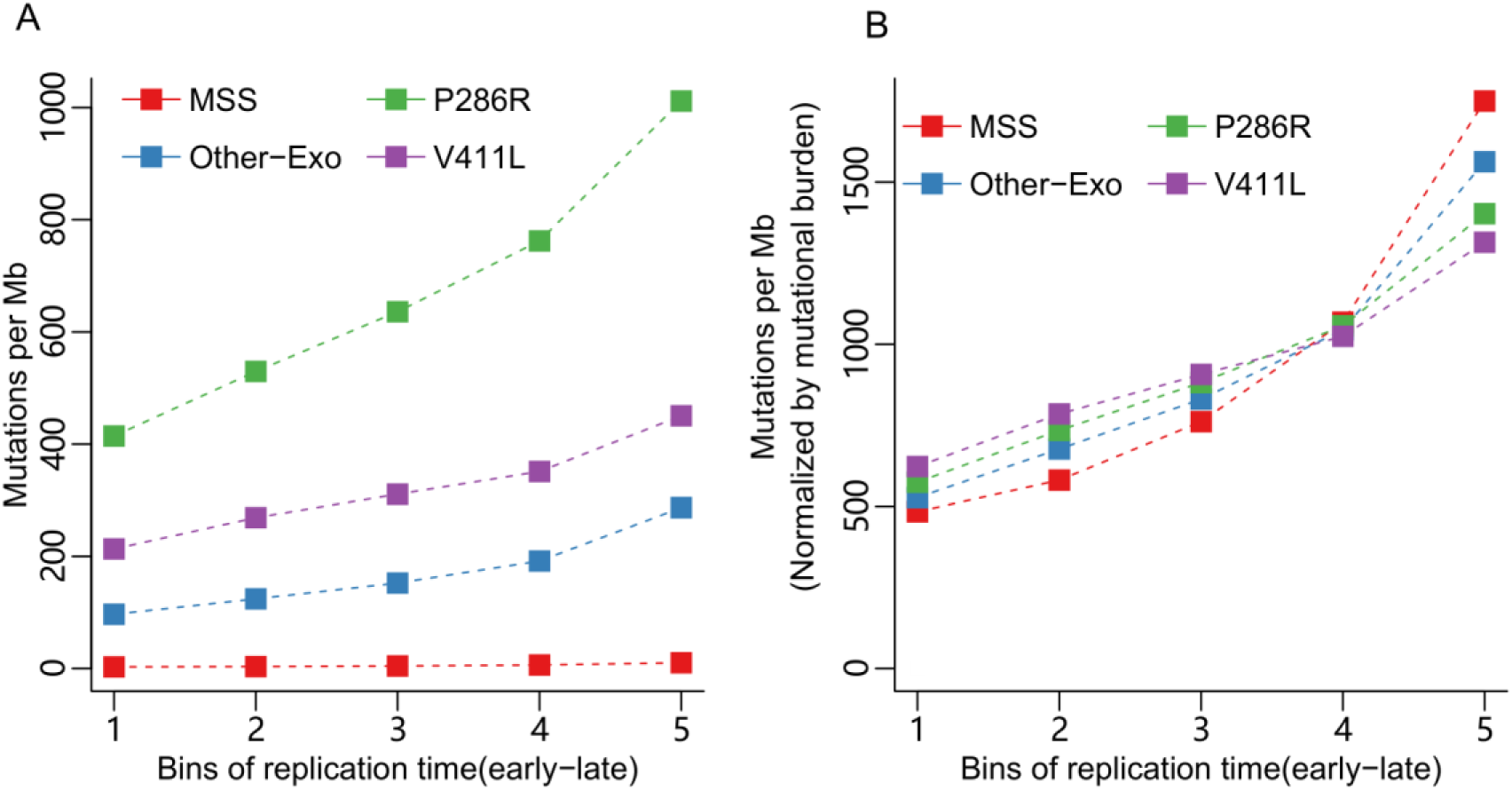
Association of mutational burden and replication timing. **(A)** DNA sequence with different replication timing was divided into 5 bins, and mutational burden was calculated in each bin ordered from early-to-late. **(B)** Mutational burden was normalized by total number of mutations in each type of mutant.

**Figure 4-figure supplement 1.**
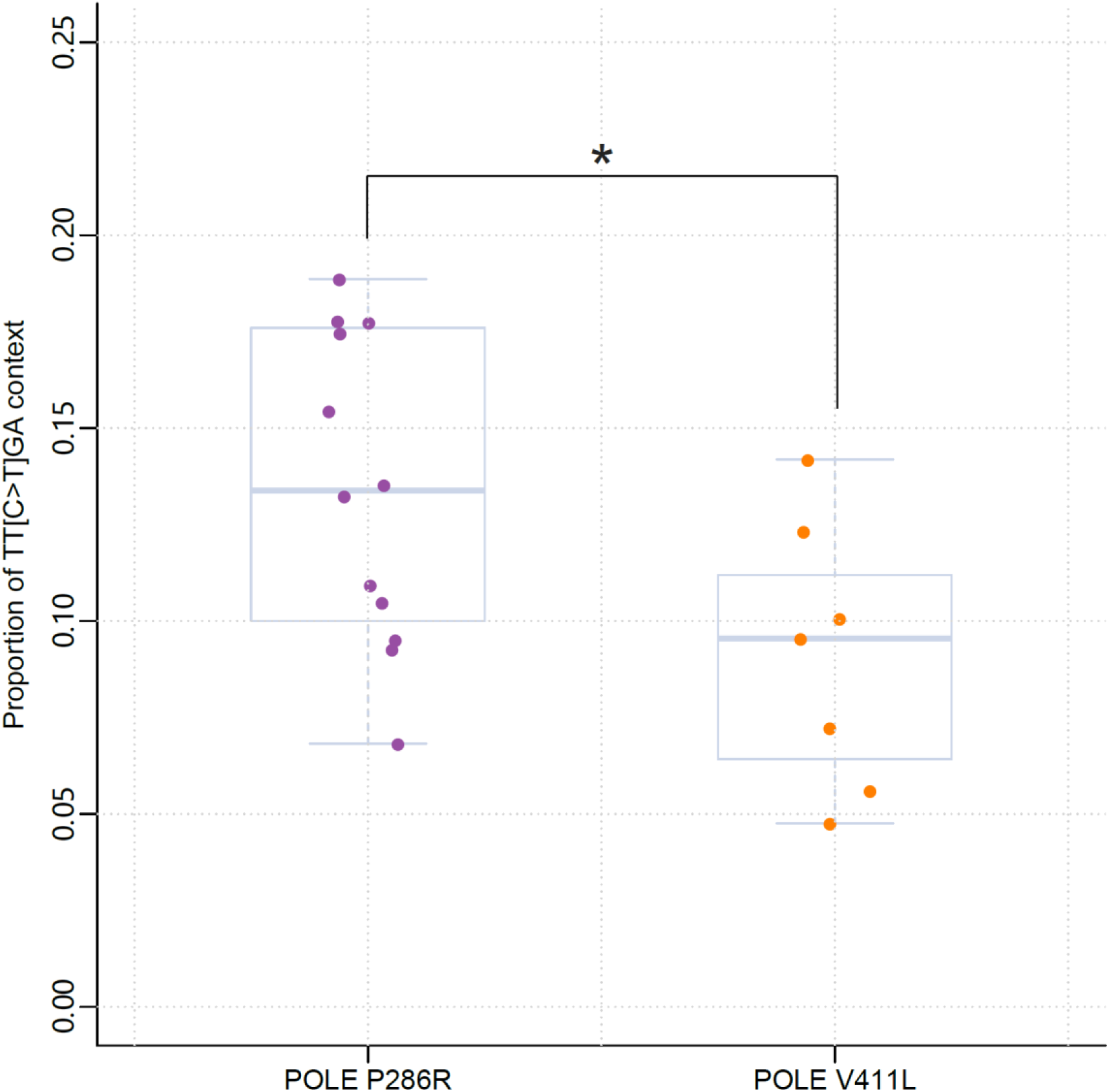
Proportion of C>T mutations in the TTCGA pentanucleotide context in POLE P286R mutants and POLE V411L mutants. * < 0.05, Student’s t-test.

## Supplementary file

The supplementary file includes the follow supplementary tables:

**Supplementary table 1**. Hotspot status across in all colorectal cancer across all cohorts

**Supplementary table 2**. Significance of enrichment of each mutation hotspot in the different POLE mutants.

**Supplementary table 3**. Extended contingency table comparing enrichment of the TP53 R213* mutation.

**Supplementary table 4**. Sample information of 53 whole genome sequencing colorectal cancer

**Supplementary table 5**. Summary of cohorts used in the study

